# Cell wall damage reveals spatial flexibility in peptidoglycan synthesis and a non-redundant role for RodA in mycobacteria

**DOI:** 10.1101/2021.10.26.465981

**Authors:** Emily S. Melzer, Takehiro Kado, Alam García-Heredia, Kuldeepkumar Ramnaresh Gupta, Xavier Meniche, Yasu S. Morita, Christopher M. Sassetti, E. Hesper Rego, M. Sloan Siegrist

**Affiliations:** Department of Microbiology, University of Massachusetts, Amherst, MA 01003, USA; Molecular and Cellular Biology Graduate Program, University of Massachusetts Amherst, MA 01003; Department of Biology, Massachusetts Institute of Technology, Cambridge, MA 02139; Department of Microbial Pathogenesis, Yale University School of Medicine, New Haven, CT 06519; Department of Microbiology and Physiological Systems, University of Massachusetts Medical School, Worcester, MA 01655

## Abstract

Cell wall peptidoglycan is a heteropolymeric mesh that protects the bacteria from internal turgor and external insults. In many rod-shaped bacteria, peptidoglycan synthesis for normal growth is achieved by two distinct pathways: the Rod complex, comprised of MreB, RodA and a cognate class B PBP, and the class A PBPs. In contrast to laterally-growing bacteria, pole-growing mycobacteria do not encode an MreB homolog and do not require SEDS protein RodA for *in vitro* growth. However, RodA contributes to survival of *Mycobacterium tuberculosis* in some infection models, suggesting that the protein could have a stress-dependent role in maintaining cell wall integrity. Under basal conditions, we find here that the subcellular distribution of RodA largely overlaps with that of the aPBP PonA1, and that both RodA and the aPBPs promote polar peptidoglycan assembly. Upon cell wall damage, RodA fortifies *M. smegmatis* against lysis and, unlike aPBPs, contributes to a shift in peptidoglycan assembly from the poles to the sidewall. Neither RodA nor PonA1 relocalize; instead, the redistribution of nascent cell wall parallels that of peptidoglycan precursor synthase MurG. Our results support a model in which mycobacteria balance polar growth and cell-wide repair via spatial flexibility in precursor synthesis and extracellular insertion.

**Importance:** Peptidoglycan synthesis is a highly successful target for antibiotics. The pathway has been extensively studied in model organisms under laboratory-optimized conditions. In natural environments, bacteria are frequently under attack. Moreover the vast majority of bacterial species are unlikely to fit a single paradigm because of differences in growth mode and/or envelope structure. Studying cell wall synthesis under non-optimal conditions and in non-standard species may improve our understanding of pathway function and suggest new inhibition strategies. *Mycobacterium smegmatis,* a relative of several notorious human and animal pathogens, has an unusual polar growth mode and multi-layered envelope. In this work we challenged *M. smegmatis* with cell wall-damaging enzymes to characterize the roles of cell wall-building enzymes when the bacterium is under attack.

## Introduction

Bacterial cell wall peptidoglycan is required for viability in most species under most conditions (1). Although peptidoglycan synthesis has been extensively studied, much of this work has been done under idealized growth conditions that do not reflect the variety of stressors found in the natural environment. Outside of the laboratory, the bacterial cell wall is under constant attack. In virtually all environments, competitors, predators, and unwilling hosts challenge bacteria with peptidoglycan-hydrolyzing enzymes (1–5). However, mechanisms to counteract cell wall damage are poorly defined. Studying peptidoglycan synthesis and remodeling under non-optimal stress conditions may lead to a better understanding of pathogenesis and ecologically-relevant pathways and interactions.

In laterally-growing, rod-shaped organisms like *Escherichia coli* and *Bacillus subtilis*, the combined activity of two distinct peptidoglycan polymerization pathways ensures cell wall integrity during normal growth as well as hostile conditions. The final, lipid-linked peptidoglycan precursor lipid II is synthesized in the inner leaflet of the plasma membrane by MurG, then flipped to the outer leaflet by MurJ (6) and inserted into the existing cell wall by the action of transglycosylases and transpeptidases. In one pathway, two dedicated enzymes work as a cognate pair, SEDS-family transglycosylase RodA (7, 8), and a monofunctional, class B penicillin-binding protein (bPBP) transpeptidase (9, 10). Along with cytoskeletal protein MreB, these proteins make up the Rod complex. This essential pathway contributes to elongation and rod-shape homeostasis by directed motion around the cell (10–13). A second, non-essential, pathway utilizes bifunctional, class A PBP (aPBPs) that perform both transglycosylation and transpeptidation (9, 14, 15), move diffusively, and are thought to maintain and repair the cell wall (16–21). Despite a growing body of evidence suggesting that aPBPs are important for stress response while the Rod complex contributes to normal growth, there are also reports that Rod complex components can sense and respond to stress (22–24).

While RodA and its cognate bPBP are more conserved than the aPBPs (25–27), they are not found in all bacterial species (28). Even when they are encoded in the genome, RodA and its bPBP are not always essential for viability nor are they always associated with MreB. For example, mycobacteria and related organisms lack MreB and do not rely on RodA for *in vitro* growth (29–33). Individual aPBPs are also largely dispensable for *in vitro* growth in this genus, with *Mycobacterium smegmatis* PonA1 a notable exception (9, 14, 15, 34). Why have these organisms retained enzymatically-redundant systems for peptidoglycan synthesis? One clue may arise from work with the human pathogen *M. tuberculosis*, where RodA and the aPBPs individually contribute to survival in immune cells, some mouse backgrounds, and in a guinea pig model (29, 35–39). These observations suggest that RodA and the aPBPs play unique roles in protecting mycobacteria from stress.

Another way that the mycobacterial cell wall differs from those of model organisms is its polar mode of elongation. Cell wall damage from external sources poses a spatial challenge to pole-growing bacteria, as it presumably is not confined to the normal sites of active peptidoglycan metabolism. We previously found that treatment with peptidoglycan-digesting enzymes lysozyme and mutanolysin causes nascent cell wall in *M. smegmatis* to shift from the poles to the sidewall (40). Here we show that *M. smegmatis* RodA and PonA1 largely overlap in localization and activity. Upon cell wall damage, peptidoglycan synthesis is redistributed from the pole to the sidewall. Neither RodA nor PonA1 relocalize in a manner that corresponds to this shift; instead, the redistribution of nascent cell wall correlates with that of peptidoglycan precursor synthase MurG. Although not essential for growth under normal laboratory conditions, RodA has a non-redundant role in damage-induced relocalization of cell wall synthesis and protects *M. smegmatis* from lysis under this condition. Our data support a model in which the location of precursor synthesis and use of specific transglycosylases can be tuned for growth or repair.

## Results

### Substantial overlap in PonA1 and RodA localization under basal conditions

In other organisms, aPBPs are hypothesized to contribute to cell wall integrity and the Rod complex, to cell wall elongation (9, 10, 16, 17, 22, 41, 42). If this division of labor is true in pole-growing mycobacteria, we hypothesized that RodA may be more polar than aPBPs like PonA1. To test this hypothesis we sought to compare the subcellular localization of fluorescent protein fusions to PonA1 and RodA. We previously confirmed the functionality of an mRFP fusion to PonA1, an essential protein in *M. smegmatis*, by allele swapping (43). Here we used the reduced cell length phenotype associated with *rodA* deletion (29) to test and confirm functionality of our RodA-mRFP construct (Fig. S1). We also showed that the fusion protein is membrane-bound, as expected, and primarily detected as full-length (Fig. S2).

Under basal conditions, we found that RodA-mRFP and, as we and others previously reported, PonA1-mRFP, are distributed along the perimeter of the mycobacterial cell (43, 44) (Figs. 1a, 1b). Neither enzyme showed clear polar enrichment but RodA-mRFP localization extended further towards the poles than PonA1-mRFP (Figs. 1b, 1c). mRFP fusions to extracellular synthetic enzymes for other layers of the mycobacterial cell envelope, including the arabinogalactan phosphotransferase Lcp1 (45), and two mycolyltransferases, *MSMEG_3580* and *MSMEG_6396* (46), showed very different patterns of localization from the peptidoglycan synthetic enzymes, indicating that the subcellular localization patterns of RodA-mRFP and PonA1-mRFP are specific (Figs. 1a, 1b). These data suggest that the cell-wide distributions of the proteins largely overlap, with RodA-mRFP slightly more polar than PonA1-mRFP.

**Figure 1:**
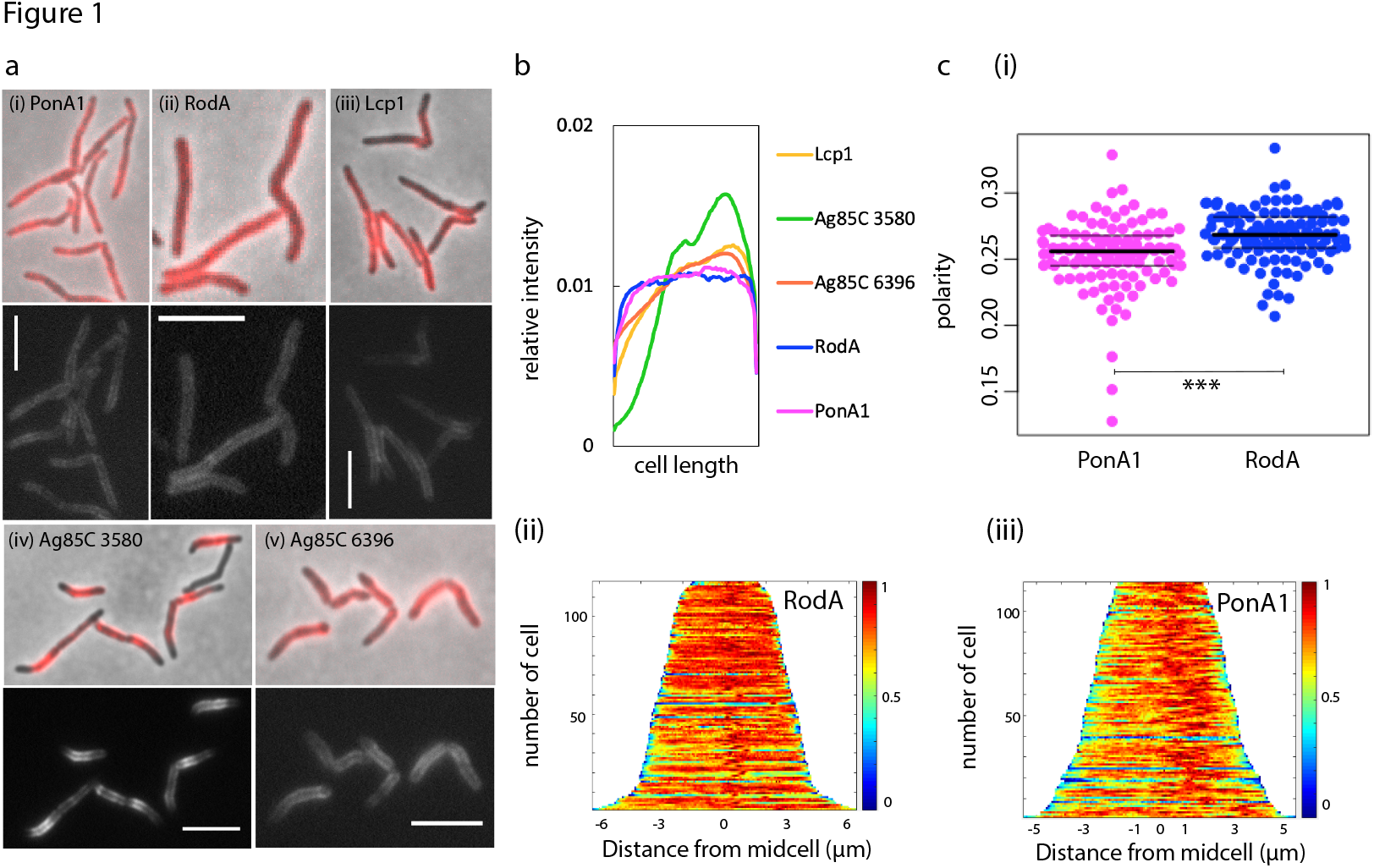
RodA and PonA1 localize around cell periphery under basal conditions. (a) Enzymes involved in (i, ii) peptidoglycan-, (iii) arabinogalactan-, and (iv, v) mycomembrane synthesis were fused to mRFP and imaged at exposure of (i) 5 s or (ii-v) 4 s. Scale bars = 5 μm. (b) Normalized fluorescence intensity profiles for mRFP tagged enzymes. 93<n<121. (c) Localization of RodA-mRFP and PonA1 compared by (i) polarity ratio calculated as the signal from 15% of the cell length on either pole, divided by signal from the entire cell. *t*-test, *p <* 0.001; (ii,iii) normalized demographs. 112<n<118.

### aPBPs and RodA both promote polar cell wall synthesis

Using a variety of metabolic labeling probes, we and others have found that active cell wall metabolism in mycobacteria occurs in asymmetric polar gradients (40, 44, 47–52). To test whether the modest difference in PonA1-mRFP and RodA-mRFP localization (Fig. 1) reflected their sites of activity, we labeled nascent cell wall using the dipeptide probe alkyne-D-alanine-D-alanine (53, 54). We previously showed that this probe incorporates into lipid-linked peptidoglycan precursors in *M. smegmatis* and therefore is a readout for cell wall biosynthesis in this species (40). To visualize aPBP activity we labeled a *rodA* knockout mutant. To enrich for RodA activity, we reduced aPBP activity by moenomycin, which specifically inhibits transglycosylation by aPBPs but not by RodA (8, 55–58).

The overall amount of cell wall labeling from Δ*rodA* or moenomycin-treated wildtype cells was reduced compared to untreated wildtype (Figs. 2a,b,d). In the absence of RodA, labeling was reduced along the sidewall and, to an even greater extent, at the poles, such that there was a net decrease in the polarity of cell wall synthesis (Fig. 2c). We observed a similar phenotype when alkyne-D-alanine-D-alanine was detected by click chemistry ligation to a different fluorophore (Fig. S3). Inhibition of transglycosylation by aPBPs also resulted in a labeling decrease along the sidewall and, to a greater extent, at the poles (Fig. 2e). These data suggest that, under basal conditions, RodA and aPBPs both contribute to polar cell wall synthesis.

**Figure 2:**
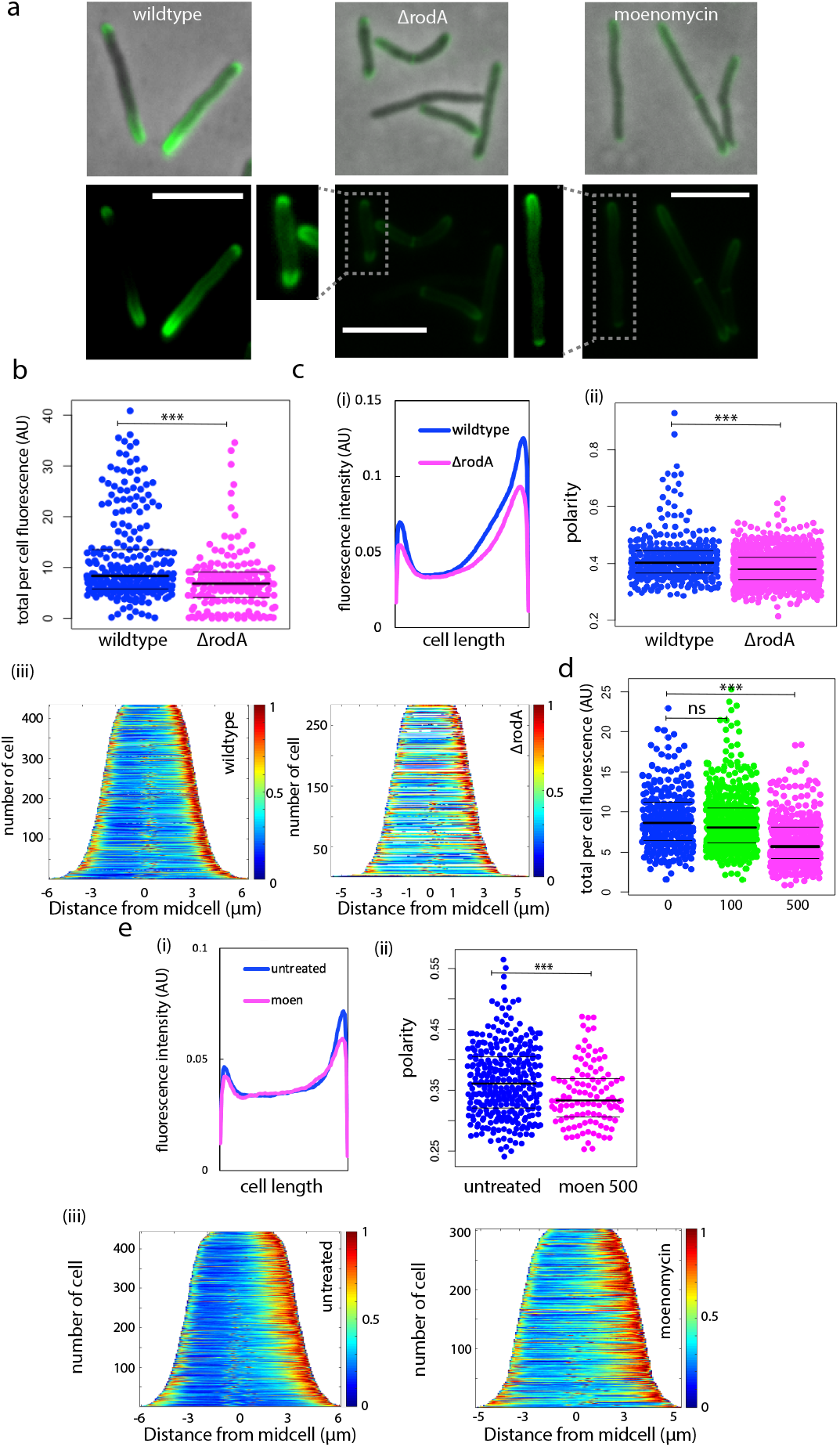
RodA and aPBPs promote polar peptidoglycan synthesis under basal conditions. (a) wildtype *M. smegmatis*, °*rodA*, and *M. smegmatis* treated with moenomycin for 30 minutes were incubated with alkyne-D-alanine-D-alanine for 10 minutes (~5-6% generation time). Probe was detected by click chemistry ligation to picolyl azide AF488. All images acquired at 1 s exposure. Dim signal from boxed cells enhanced for visibility. Scale bars = 5 μm (b) Total fluorescence per cell. *t*-test, *p* < 0.001 (c) Fluorescence signal localization of wildtype *M. smegmatis* and °*rodA* represented by (i) raw fluorescence intensity profile; (ii) polarity ratio. *t*-test, *p* < 0.001 (iii) demographs. 168<n<816. Representative of three independent experiments (d) Total fluorescence per cell following 30 minutes of 0, 100, or 500 μg/mL moenomycin. Significance determined using analysis of variance (ANOVA) followed by a Tukey post hoc test to conduct pairwise comparisons. ***, *p* < 0.001 (e) Fluorescence signal localization from wildtype cells untreated and treated with 100 or 500 μg/mL moenomycin for 30 minutes represented by (i) raw fluorescence intensity profile; (ii) polarity ratio, *t*-test, ***, *p* < 0.005; (iii) demographs. 285<n<433.

### Mutanolysin/lysozyme-mediated cell wall damage occurs along the cell periphery

We previously showed that when *M. smegmatis* cells are treated with a combination of lysozyme and mutanolysin, nascent peptidoglycan redistributes from the poles to the sidewall (40). These enzymes are glycoside hydrolases and break the linkages that connect neighbouring glycans *N*-acetylglucosamine and *N*-acetylmuramic acid in the peptidoglycan backbone (Fig. S4) (3–5). We hypothesized that cell wall assembly can shift to places where the cell wall is damaged, presumably for repair. Implicit in this hypothesis is the assumption that enzymes added to the bacterial growth medium attack the cell wall indiscriminately, *i.e.* without preference for poles vs. sidewall. In support, a previous scanning electron microscopy study showed that lysozyme-associated surface irregularities occur along the entire *M. smegmatis* cell periphery (59). We also observed that wildtype *M. smegmatis* treated with lysozyme and mutanolysin often has bumps around the cell, which we interpret as areas of weakened peptidoglycan (Fig. S4), and that loss of fluorescently-labeled cell wall occurs along the sidewall (Fig. S4).

### MurG relocalizes to the sidewall in response to cell wall damage but PonA1, RodA and DivIVA do not

We next considered what element(s) of cell wall assembly machinery might be responsible for redistributing peptidoglycan synthesis from sites of polar growth to sites of sidewall damage. After treatment with mutanolysin/lysozyme, we initially found that the localization of RodA-mRFP, and to a lesser extent, PonA1-mRFP, became more polar (Fig. S5). This was unexpected since nascent cell wall localization shifted in the opposite direction, *i.e*. became less polar (40) (Fig. 4a). However when we stained enzyme-treated cells with SYTOX Green, a dye that preferentially labels dead bacteria, we did not observe any viable cells with relocalized RodA-mRFP (Fig. S6). While we do not yet understand this phenotype, quantification of RodA-mRFP fluorescence from SYTOX Green-excluded cells suggests that cell wall damage likely does not change the localization of extracellular synthesis proteins (Fig. S6).

**Figure 3:**
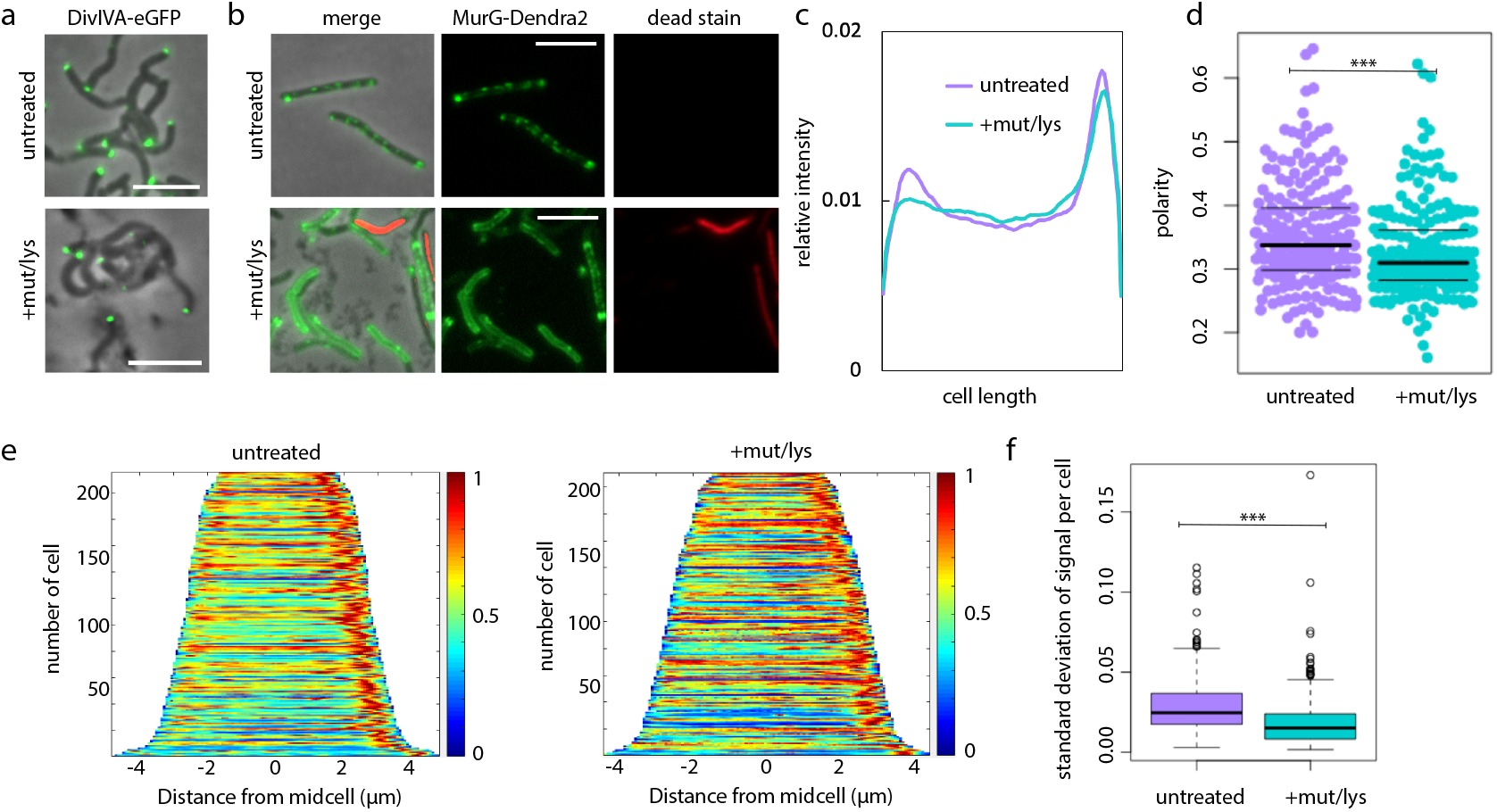
MurG relocalizes upon cell wall damage. (a) Cells expressing the DivIVA-eGFP (50) were treated +/− mutanolysin/lysozyme. Scale bars = 5 μm (b) Cells expressing the functional fusion MurG-Dendra2 (43) were treated +/− mutanolysin/lysozyme, then stained with propidium iodide for detection of dead cells. 3 s exposure for green channel, 500 ms exposure for red channel. Scale bars = 5 μm. (c) Normalized MurG-Dendra2 fluorescence intensity profiles (d) Polarity ratio of MurG-Dendra2 signal. *t*-test, *p* < 0.001. (e) MurG-Dendra2 demographs. (f) Standard deviation calculated for 100 fluorescence values per cell in untreated and treated cells. *t*-test, *p* < 0.001. 210<n<215.

**Figure 4:**
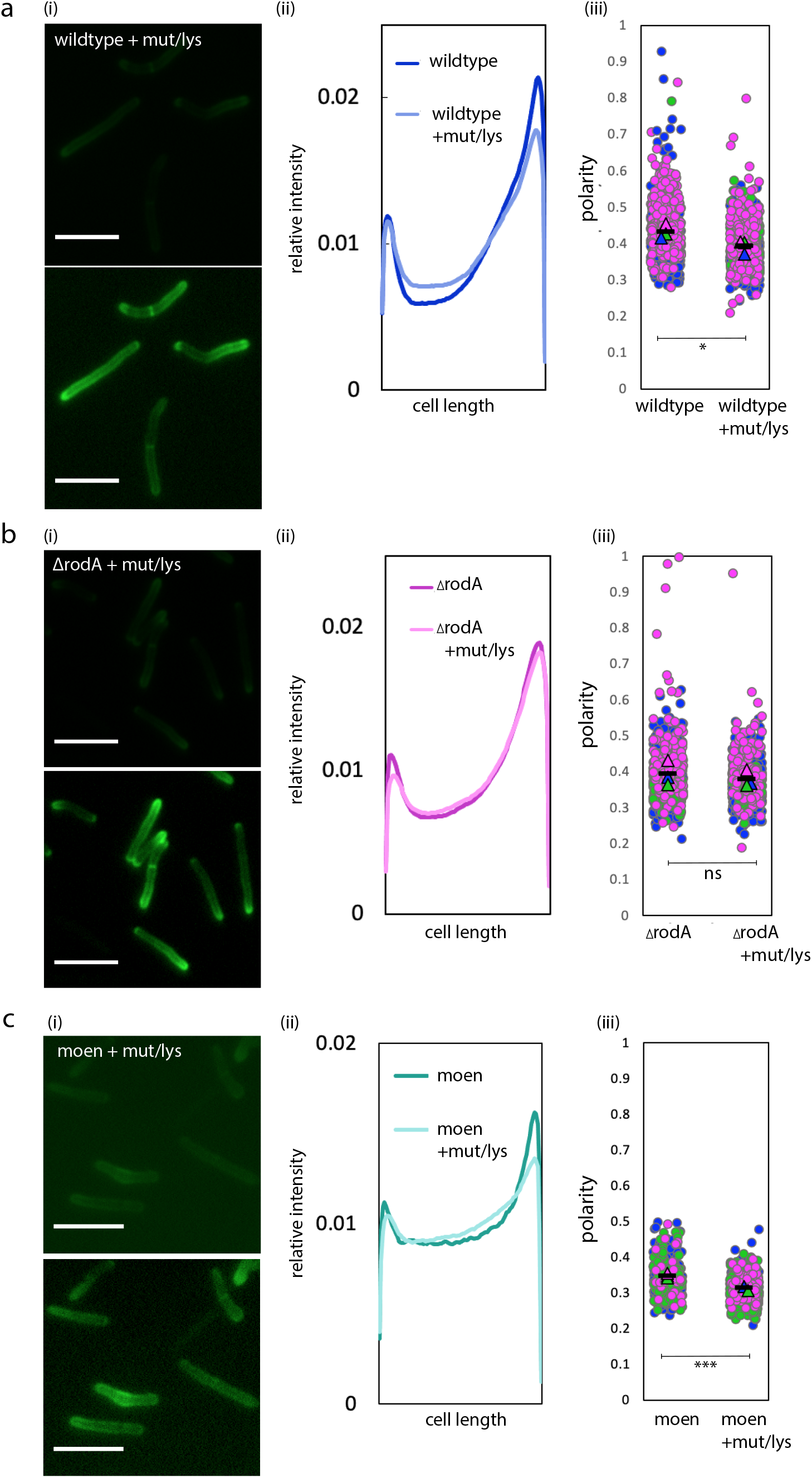
RodA contributes to damage-induced relocalization of peptidoglycan assembly. At the end of lysozyme/mutanolysin treatment, nascent cell wall in wild-type or Δ*rodA* (b) or moenomycin treated cells (c) was labeled with alkyne-D-alanine-D-alanine as in Fig. 2a and (i) imaged at 1s exposure (bottom panel is the same image with enhanced signal for visualization), compare to untreated in Fig. 2a (ii) Normalized fluorescence intensity profiles comparing relative signal from untreated and treated cells. (iii) Super plots of cell wall labeling polarity ratio (signal from both poles divided by total cell fluorescence) from 3 independent experiments, each color represents an experiment. *t*-test in (a), p<0.05, (b), *p* > 0.1, (c), *p*<0.01 102<n<826. Scale bars = 5 μm.

In contrast to well-studied, rod-shaped species, mycobacteria coordinate cell wall synthesis via the cytoskeletal protein DivIVA (Wag31), rather than MreB (50, 60–62). We wondered whether DivIVA might also organize cell wall synthesis in response to sidewall damage. However the location of the functional fluorescent protein fusion DivIVA-eGFP (40, 50, 62), like those of PonA1 and RodA, did not change upon mutanolysin/lysozyme treatment (Fig. 3a).

We next turned our attention to the source of peptidoglycan precursor substrates for the aPBPs and RodA. Synthesis of the final precursor lipid II is carried out by MurG. Accordingly we treated cells expressing the functional fluorescent protein fusion MurG-Dendra2 (43) with mutanolysin/lysozyme. Imaging of cells before and after treatment revealed that MurG-Dendra2 signal transitions from a predominantly sub-polar and patchy signal to a more uniform signal that often surrounds the entire periphery of the cell (Fig. 3b). Because relocalization of RodA-mRFP was only observed in dead cells, we stained both untreated and treated cells with propidium iodide, another dye that preferentially labels dead bacteria. After omitting propidium iodide-stained cells from our analysis, fluorescence quantitation supported our observation that MurG-Dendra2 relocalizes away from the polar region upon cell wall damage (Figs. 3c-e) and that it becomes less patchy (Fig. 3f).

Taken together, our data indicate that MurG, but not PonA1, RodA or DivIVA, relocalizes upon cell wall damage.

### RodA, but not aPBPs contributes to redistribution of peptidoglycan synthesis upon cell wall damage

The distribution of MurG, and therefore lipid II, likely contributes to spatial flexibility in peptidoglycan assembly. The location of RodA (and likely, PonA1) did not obviously change upon cell wall damage (Fig. S5, S6). However, given the basal, cell-wide distribution of both transglycosylases, we asked if they might contribute to damage-induced pole-to-sidewall redistribution of nascent peptidoglycan. We first reproduced the cell wall labeling phenotype in mutanolysin/lysozyme-treated wildtype *M. smegmatis* (40) and showed that there was a decrease in nascent peptidoglycan polarity (Fig. 4a). By contrast, we found that mutanolysin/lysozyme treatment of *M. smegmatis* lacking RodA did not change the polarity of nascent peptidoglycan (Fig. 4b, Fig. S3). However, pretreating wildtype cells with the aPBP-transglycosylation inhibitor moenomycin did not prevent pole-to-sidewall damage response (Fig. 4c). These data suggest that RodA, but not aPBPs, contributes to stress-induced spatial flexibility in peptidoglycan synthesis.

### A non-redundant role for RodA in protection against cell wall damage

The contribution of RodA_Mtb_ to *M. tuberculosis* survival in different *in vivo* models (29, 35–37, 39) and the requirement for RodA_Msm_ in the damage-induced sidewall shift in *M. smegmatis* peptidoglycan synthesis (Fig. 4) suggested that the enzyme could play a non-redundant role in protecting against cell wall stress. To test, we challenged wild-type and Δ*rodA M. smegmatis* cultures in the presence or absence of moenomycin with mutanolysin/lysozyme and plated spot dilutions (Fig. S7). For reasons that we do not yet understand, moenomycin treatment appeared to reduce the sensitivity of *M. smegmatis* to cell wall damage. Because the spot dilutions did not have the resolution to test whether there was a difference between wildtype and Δ*rodA*, we performed full-plate colony-forming unit (CFU) assays and microscopy at several time points. Upon addition of mutanolysin/lysozyme to the growth medium, wildtype *M. smegmatis* grew for 2 hours prior to lysing (Fig. 5a) albeit more slowly than untreated cells (Fig. 5b). By contrast Δ*rodA* immediately began to lyse, a phenotype evident both by CFUs and by microscopy (Fig. 5a, Fig. S4). Δ*rodA* lysis was complemented by expression of *rodA-* mRFP (Fig. 5b). Expression of *rodA-*mRFP in a wildtype background, moreover, enhanced survival compared to wildtype alone (Fig. 5b). Thus while RodA is dispensable for normal growth (29) (Fig. S8), it plays a non-redundant role in protection from cell wall damage.

## Discussion

We have previously shown that upon cell wall insult, peptidoglycan assembly in *M. smegmatis* relocalizes from the growing poles to the non-growing sidewall (40). Given that bacteria are likely to encounter such insults frequently in their natural habitats, we sought to better understand the factors that drive relocalization. We focused on the roles of two peptidoglycan transglycosylases, the aPBP PonA1 and SEDS family transglycosylase RodA.

**Figure 5:**
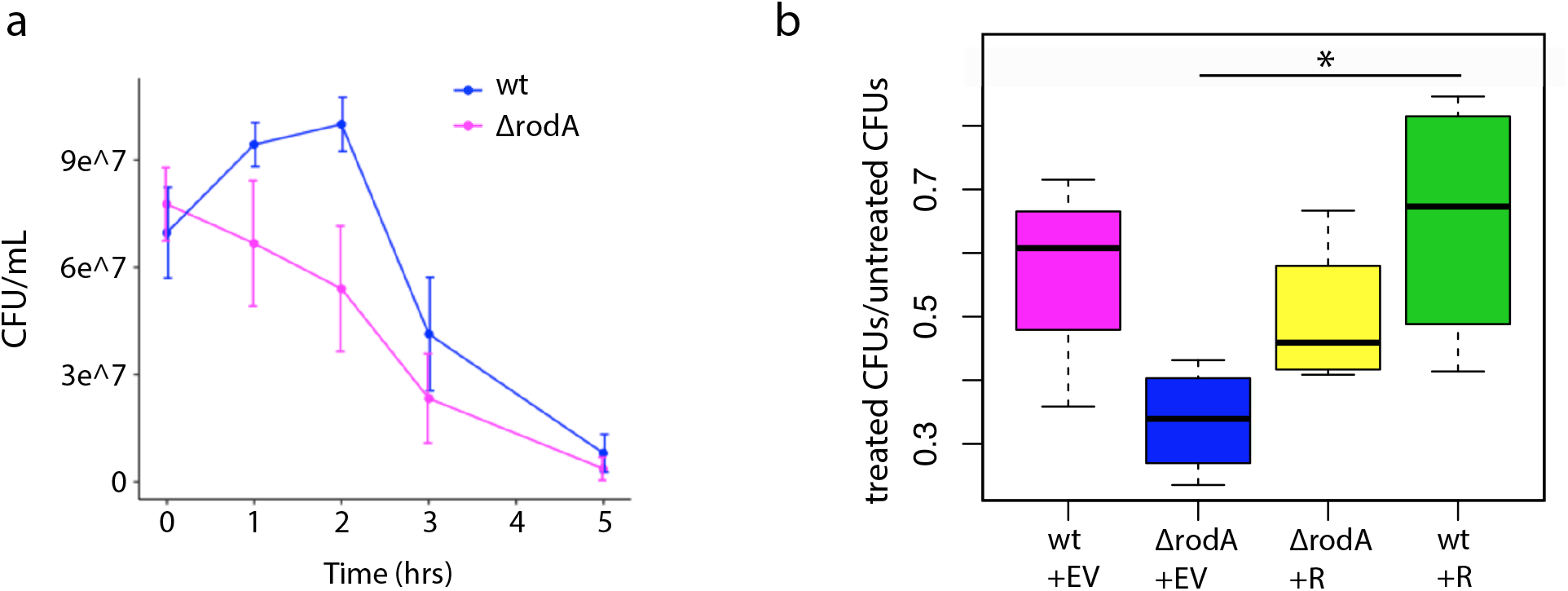
RodA protects against cell wall damage. (a) Colony forming units (CFU) over time for wildtype and Δ*rodA* cells +/− mutanolysin/lysozyme. *n* = 3. Compare to Figure S8 (b) Ratios of treated/untreated CFUs after 2 hours of mutanolysin/lysozyme, comparing wildtype + empty vector (EV), Δ*rodA +* empty vector, Δ*rodA + rodA-mRFP* (R), wildtype + *rodA-mRFP*. While cultures were normalized to OD_600_ = 0.05 prior to treatment, the uncomplemented mutant and RodA overexpression strains consistently had higher and lower CFU than wildtype at t = 0, respectively. This is likely due to the differences in cell length between the four strains (29) (Fig. S1). To better highlight the effects of mutanolysin/lysozyme we calculated the fold change in CFU between treated and untreated cells after two hours. *n* = 4 independent experiments, two of which done in triplicate. Significance determined using analysis of variance (ANOVA) followed by a Tukey post hoc test to conduct pairwise comparisons. *, *p* < 0.05, only significant relationship portrayed.

In laterally-growing, rod-shaped bacteria, the emerging narrative is that RodA lays the template for elongation and aPBPs fill in the gaps for maintenance and repair (10, 16–19, 21). Unlike the organisms in which this model has been tested, pole-growing bacteria like members of the Mycobacteriales and Hyphomicrobiales lack the cytoskeletal protein MreB and either lack or do not require RodA for viability or shape (28, 29). If the division of labor that has been reported in laterally-growing bacilli were employed by mycobacteria, we would predict that localization and activity of RodA would be enriched at the poles, while localization and activity of aPBPs like PonA1 would be distributed around the cell periphery. This is reminiscent of the model for transpeptidation in mycobacterial peptidoglycan, where the D,D-transpeptidases that catalyze 4,3-crosslinks associated with lipid II insertion are likely to be enriched at the poles and the L,D-transpeptidases that catalyze 3,3-crosslinks associated with cell wall maturation localize along the cell periphery (40, 44, 63, 64). Instead we observed that the distributions of functional fluorescent protein fusions to RodA and PonA1 are similar, as are their enzymatic activities (Figs. 1-2). As recent findings in pole-growing *Corynebacterium glutamicum* (31) also suggest, the division of labor for *M. smegmatis* peptidoglycan polymerases under basal conditions is subtle.

While RodA and aPBPs make similar contributions to polar growth, our data suggest that their roles diverge upon cell wall damage (Fig. 5). Specifically, RodA plays a non-redundant role in damage-associated pole-to-sidewall redistribution of peptidoglycan synthesis, which is concomitant with a similar redistribution of the lipid II synthase MurG (Fig. 3). It is not yet clear if the change in substrate availability is the cause, consequence or simply occurs in parallel with the change in transglycosylase usage. In the future, localization of lipid II flippase MurJ—which bridges lipid II synthesis in the inner leaflet and lipid II insertion in the outer leaflet—may help us to distinguish between these models. In *Staphylococcus aureus*, MurJ recruitment redirects peptidoglycan synthesis from the cell periphery, for expansion, to midcell, for division (65).

Our data suggest that RodA promotes a pole-to-sidewall shift in peptidoglycan synthesis (Fig. 4) and survival during cell wall damage (Fig. 5). The non-redundant role(s) for RodA in resistance to lysis (Fig. 5) is consistent with at least two models. In the first, RodA provides on-demand repair of cell wall damage. A similar stress-specific, peptidoglycan-building role for RodA has been suggested in *Listeria monocytogenes* (24), where absence of a RodA homolog also sensitizes bacteria to cell wall damage (66). Loss of damage-induced sidewall shift supports this type of active role for RodA in mycobacteria. However if true, this would be in contrast to the repair function of aPBPs, rather than RodA, in other organisms (16–19, 21). Thus a second model to explain the lysis phenotype of Δ*rodA* is that RodA builds a cell wall with an architecture that is inherently more resistant to damage or that is more amenable to repair.

The utility of two pathways for the same enzymatic reaction is not clear under laboratory-optimized growth conditions. By studying the requirements for peptidoglycan synthesis during cell wall damage, we have uncovered spatial flexibility in precursor synthesis and extracellular insertion, and a non-redundant role for RodA in protection (Fig. 6). These factors may enable mycobacteria to balance polar growth with cell-wide repair in the host and soil environments.

**Figure 6:**
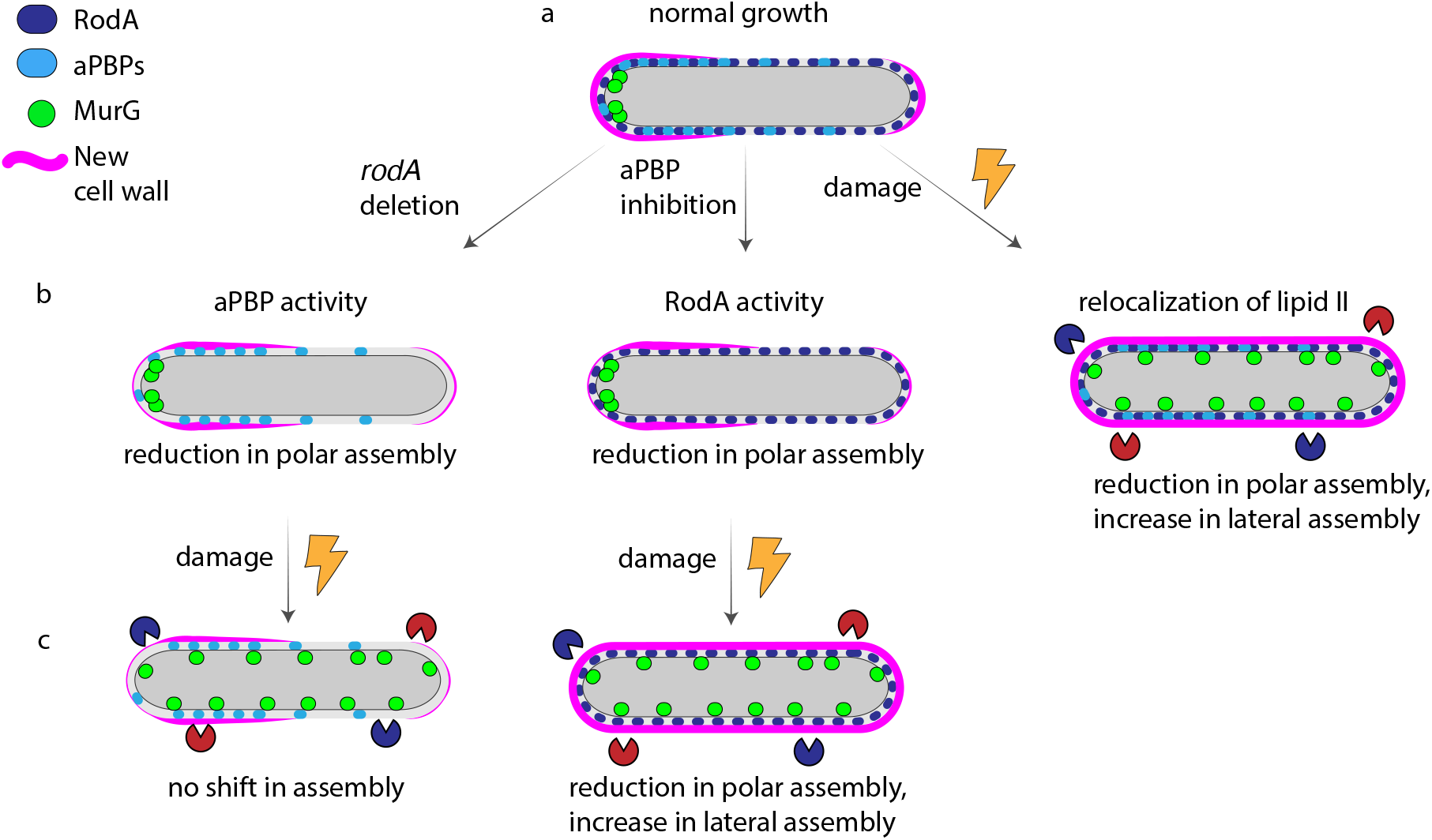
Spatial flexibility model for peptidoglycan assembly. (a) under basal conditions PonA1 and RodA overlap substantially but not completely; new cell wall is assembled asymmetrically and enriched at the poles. (b, left and middle) upon *rodA* deletion and aPBP inhibition new cell wall assembly is disproportionally reduced at the poles. (b, right) upon cell wall damage MurG redistributes from the poles to the cell periphery, as does new cell wall. (c, left) in the absence of RodA cell wall assembly is not shifted to the lateral cell under stress. (c, right) when aPBPs are inhibited cell wall assembly redistributes upon damage.

## Materials and Methods

### Strain construction

Genes of interest were amplified from *M. smegmatis* mc^2^155 genomic DNA. *mRFP* was amplified from a pL5pTetO plasmid, with primers in Table S1. Backbone plasmid pL5pTetO was linearized by PCR. 15 ng of plasmid backbone, 20 ng of gene of interest, and 20 ng of *mRFP* PCR products were incubated with Gibson master mix (New England Biolabs, #E2611S) at 50°C for 1 hour. 5 μL of Gibson product was transformed into 50 μL XL1-Blue *E. coli* competent cells by heat shock. Transformants on 50 μg/mL kanamycin plates were confirmed by colony PCR and sequencing, and then electroporated into *M. smegmatis* mc^2^155 or into °*rodA*.

### Cell wall damage

Wildtype or *ΔrodA* cells were grown to stationary phase, then back-diluted and allowed overnight growth to log phase (OD_600_ = 0.5-0.8). Lysozyme (Sigma-Aldrich, #L6876) was freshly resuspended in PBS, filter-sterilized, and added to cultures at a final concentration of 500 μg/mL. Mutanolysin (Sigma-Aldrich #M9901) was added simultaneously at a final concentration of 500 U/mL. Cultures were incubated at 37°C shaking at 300 rpm for 1 hour.

### Peptidoglycan labeling

Peptidoglycan precursor probe alkyne-D-alanine-D-alanine (2 mM; custom synthesized by WuXi AppTech), was added to cultures for the final 10°minutes of incubation. Cells were washed three times in cold PBS and fixed in 2% formaldehyde for 10 minutes. Cells were washed in PBS then subjected to CuAAC with picolyl azide-AF488 or picolyl azide-TAMRA (Click Chemistry Tools) as described (67, 68).

### CFUs and growth curves

Wildtype + pL5pTetO, wildtype + pL5pTetO-*rodA-mRFP*, ΔrodA + pL5pTetO, and ΔrodA + pL5pTetO-*rodA-mRFP* cells were grown to stationary phase, then back-diluted and allowed to grow overnight to log phase (OD_600_ = 0.5-0.8). Cultures were back-diluted once more to OD_600_ = 0.05. Lysozyme and mutanolysin were added as above. Triplicate cultures were incubated at 37°C shaking at 150 rpm for 5 hours. Aliquots were periodically plated for CFUs.

### Moenomycin treatment

Wildtype cells were grown to stationary phase, then back-diluted and allowed to grow overnight to log phase (OD_600_ = 0.5-0.8). Moenomycin (Cayman Chemicals, #15506) was added at concentrations described in text. Cultures were incubated at 37°C shaking at 400 rpm for 30 minutes in Benchmark Scientific MultiTherm Shaker H5000-H.

### Viability staining

Staining was calibrated using untreated cells as live control and 70% isopropanol-treated cells as dead control. Following 90 minute treatment with lysozyme and mutanolysin, cells expressing RodA-mRFP were washed with HEPES-buffered saline (HBS) twice, and resuspended in HBS + Sytox Green (Fisher Scientific, #S7020) at a final concentration of 2.5 μM. For cells expressing MurG-Dendra2, propidium iodide was added to a final concentration of 4 μM. Cells were then incubated in the dark for an additional 30 minutes and imaged immediately.

### Imaging

Cells were placed on pads made of 1% agarose in water. Images were acquired on Nikon Eclipse E600 at exposure times detailed in main text.

### Image analysis

Cell outlines were traced using Oufti (69). Demographs were generated using tools built into the program. Intensity profiles of non-septating labeled cells only were generated using MATLAB code described in (40). Polarity ratios were calculated by combining signal values for 15% of the cell length on either pole and dividing the sum by total cell fluorescence. Beeswarm plots and boxplots generated on R studio. Super plots were generated as described in (70).

### Membrane fractionation and western blotting

Wildtype *M. smegmatis* and cells expressing RodA-mRFP were grown to OD_600_ ~0.6 and lysed by nitrogen cavitation. Lysates were separated into cytoplasm and membrane fractions by ultracentrifugation at 35,000 rpm for 2 hours. Protein concentration was adjusted to 560 μg/mL. Cell lysate or fractionated samples were separated by SDS–PAGE on a 4-20% polyacrylamide gel and transferred to a PVDF membrane. The membrane was blocked with 3% skim milk in PBS + 0.05% Tween-80 (PBST) and then incubated overnight with primary monoclonal mouse anti-RFP. Antibodies were detected with appropriate secondary antibodies conjugated to horseradish peroxidase (GE Healthcare, Chicago, IL). Membranes were rinsed in PBS + 0.05% Tween-20 before visualization.

## Supplementary data

**Figure S1:**
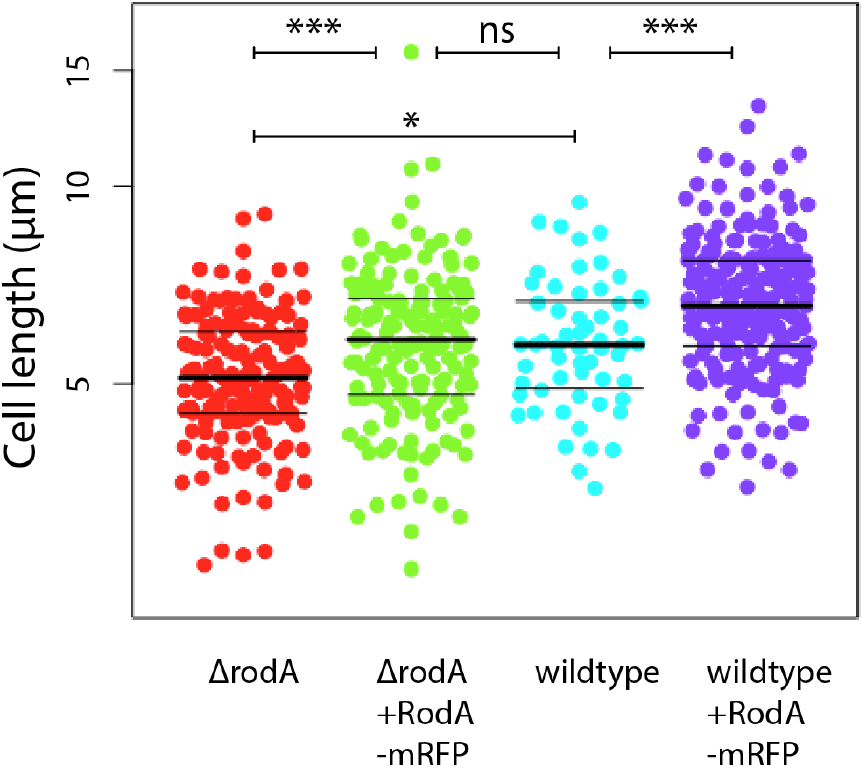
RodA-mRFP fusion complements Δ*rodA*. Phase images of ΔrodA + empty vector, ΔrodA + *rodA-mRFP*, wildtype + empty vector, and wildtype + *rodA-mRFP* were acquired and cell lengths were quantified using Oufti (69), MATLAB, and visualized using R Studio (58<n<204). Significance determined using analysis of variance (ANOVA) followed by a Tukey post hoc test to conduct pairwise comparisons. ns, not significant; *, p<0.05; **, p<0.01; ***, p<0.005.

**Figure S2:**
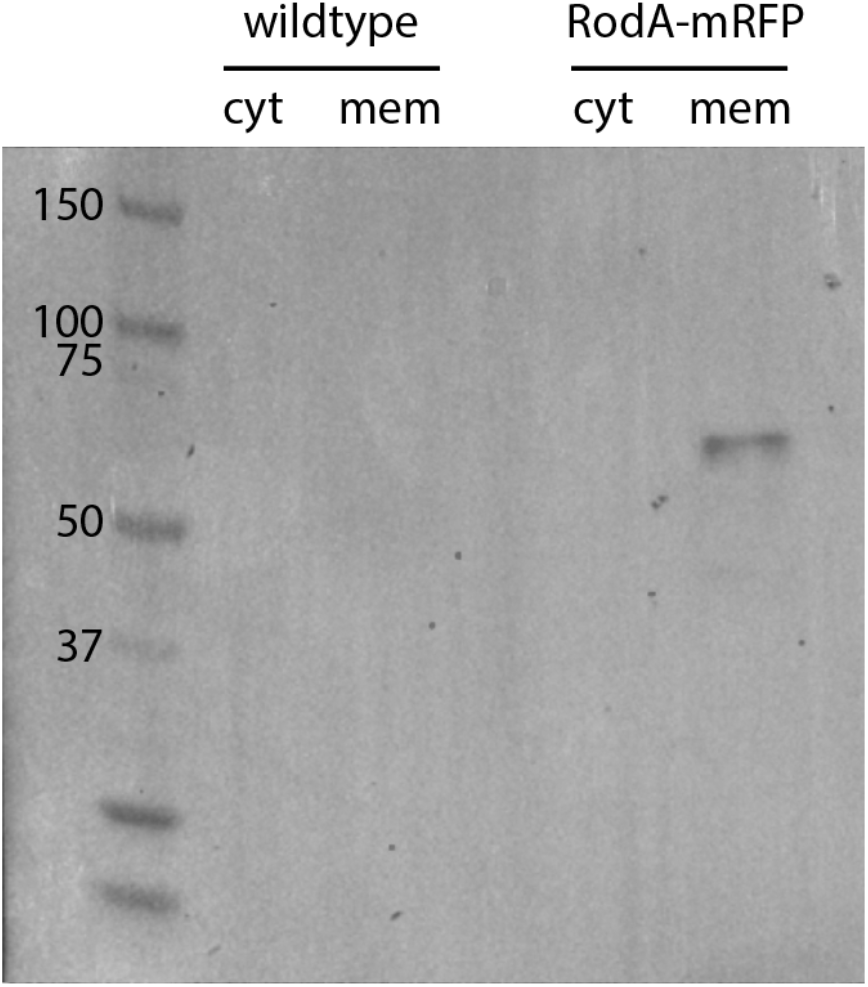
RodA-mRFP fusion localizes to the plasma membrane with minimal degradation. Lysates from *M. smegmatis* +/− *rodA-mRFP* were separated into cytoplasmic (cyt) and membrane (mem) fractions by ultracentrifugation and immunoblotted with anti-RFP antibodies. Protein concentration normalized.

**Figure S3:**
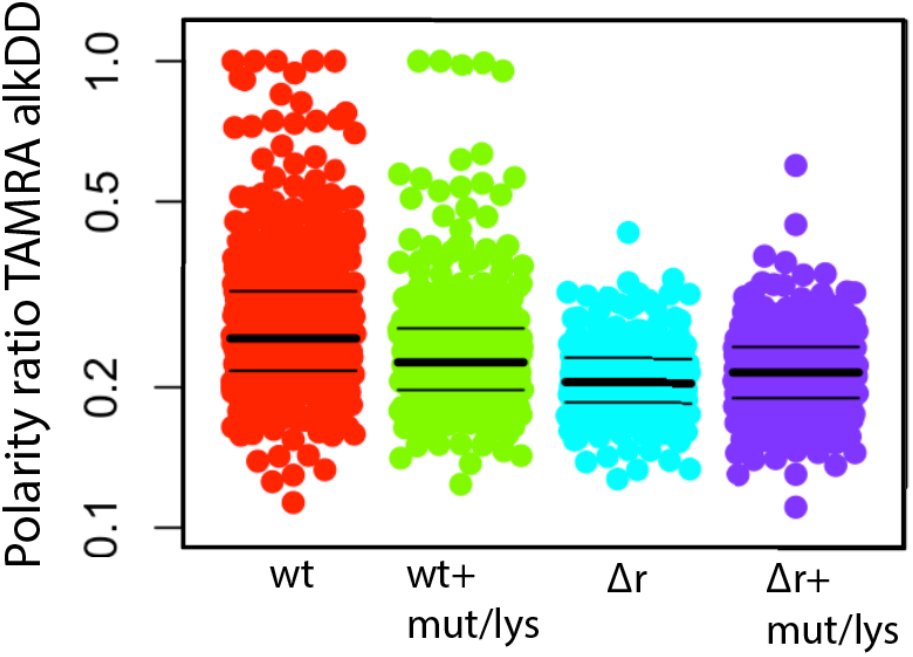
Polarity of cell wall synthesis detected with TAMRA. Polarity ratio of cell wall labeling (bright pole signal/total cell fluorescence) in wildtype and Δ*rodA* +/− mutanolysin/lysozyme. Nascent peptidoglycan labeled as in Fig. 2a except that click chemistry detection was with picolyl azide-TAMRA.

**Figure S4:**
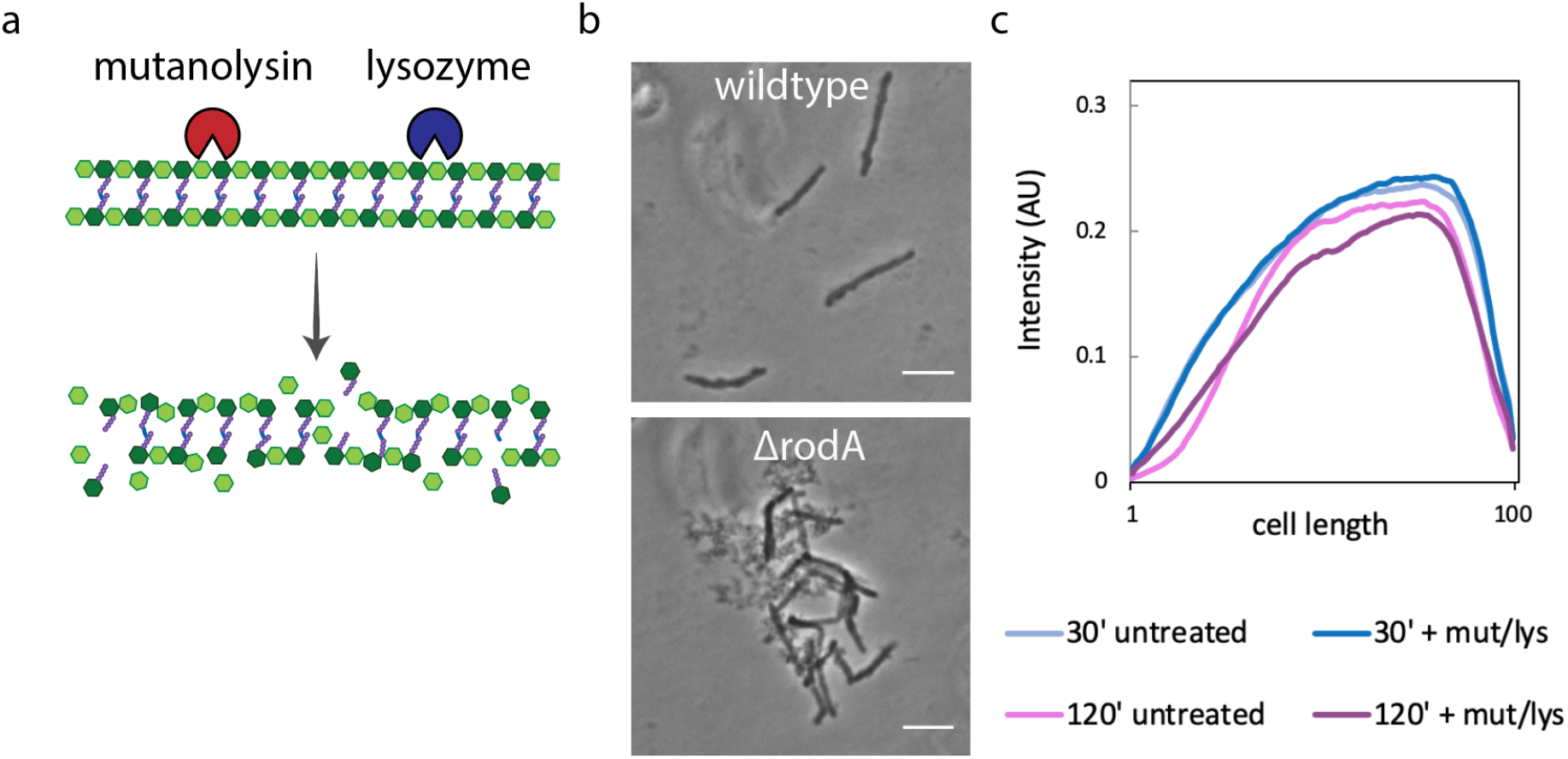
Mutanolysin/lysozyme treatment leads to cell-wide damage. (a) Muramidases mutanolysin and lysozyme break the linkages between neighbouring glycans *N*-acetylglucosamine and *N*-acetylmuramic acid in the peptidoglycan backbone (b) Phase contrast images of wildtype or Δ*rodA M. smegmatis* +/− mutanolysin/lysozyme. Scale bars = 5 μm. (c) *M. smegmatis* was labeled with 1μM RADA, a D-amino acid monopeptide that we previously showed incorporates into peptidoglycan via L,D-transpeptidases (40) and coats the cell wall evenly after overnight incubation with a low concentration (62). After washing, the culture was treated with mutanolysin/lysozyme and loss of fluorescence along the length of the cells was quantitated. As expected, after 2 hours labeling loss was observed at the poles in untreated cells. Loss of fluorescence at the poles of lysozyme/mutanolysin-treated *M. smegmatis* was not as pronounced, consistent with its slow growth in the presence of the enzymes (Figure 5a). At this time point, sidewall loss of fluorescence was greatly enhanced with enzyme treatment. Although mycobacterial growth precludes interpretation of cell wall loss at the poles, these data suggest that lysozyme/mutanolysin-mediated cell wall loss occurs along the *M. smegmatis* sidewall. Signal not normalized. 58<n<102.

**Figure S5:**
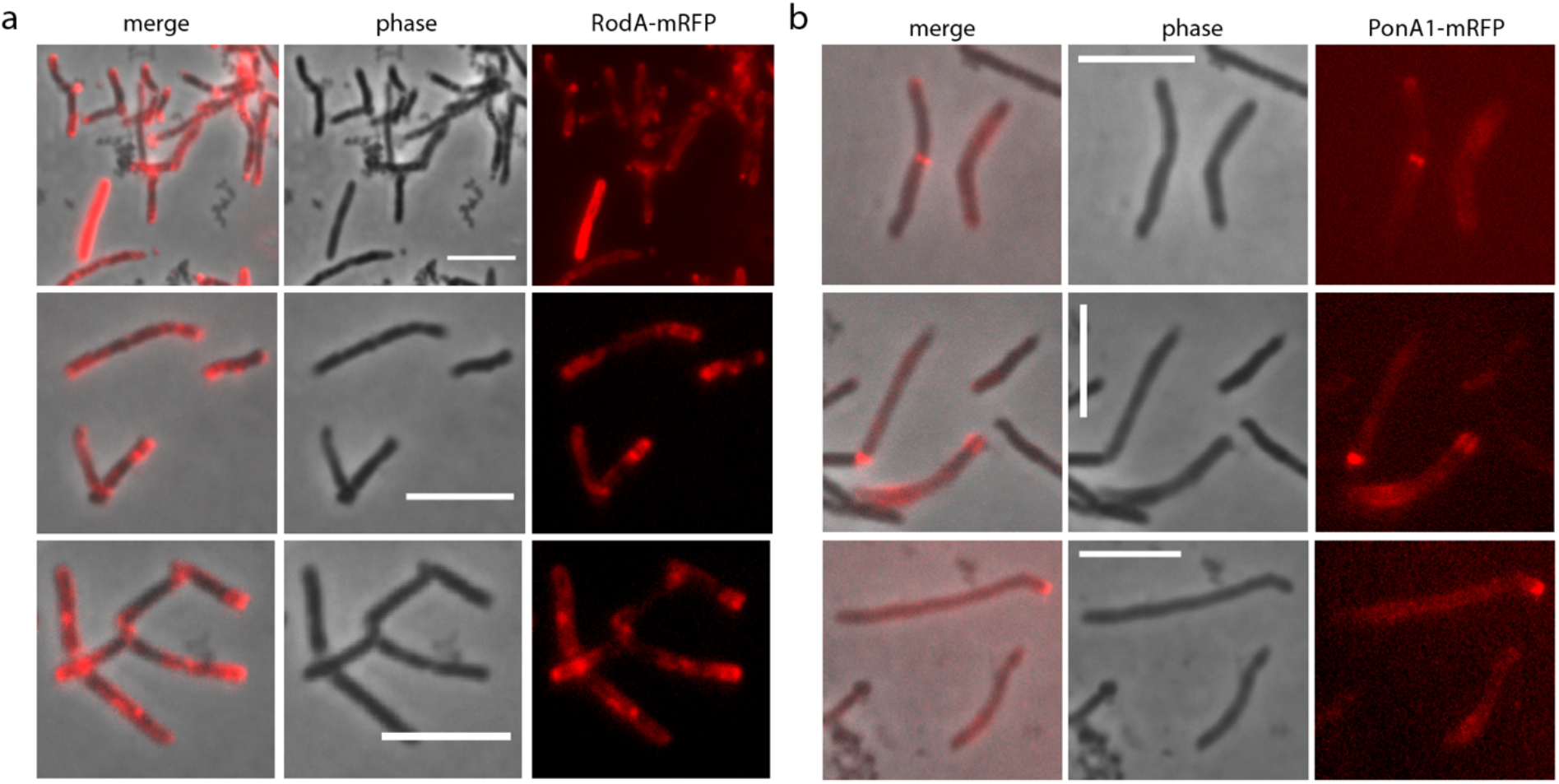
RodA-mRFP and PonA1-mRFP location in cells treated with lysozyme/mutanolysin. Representative images of (a) RodA-mRFP and (b) PonA1-mRFP imaged following mutanolysin/lysozyme treatment. Compare to Figure 1a. Scale bars = 5 μm.

**Figure S6:**
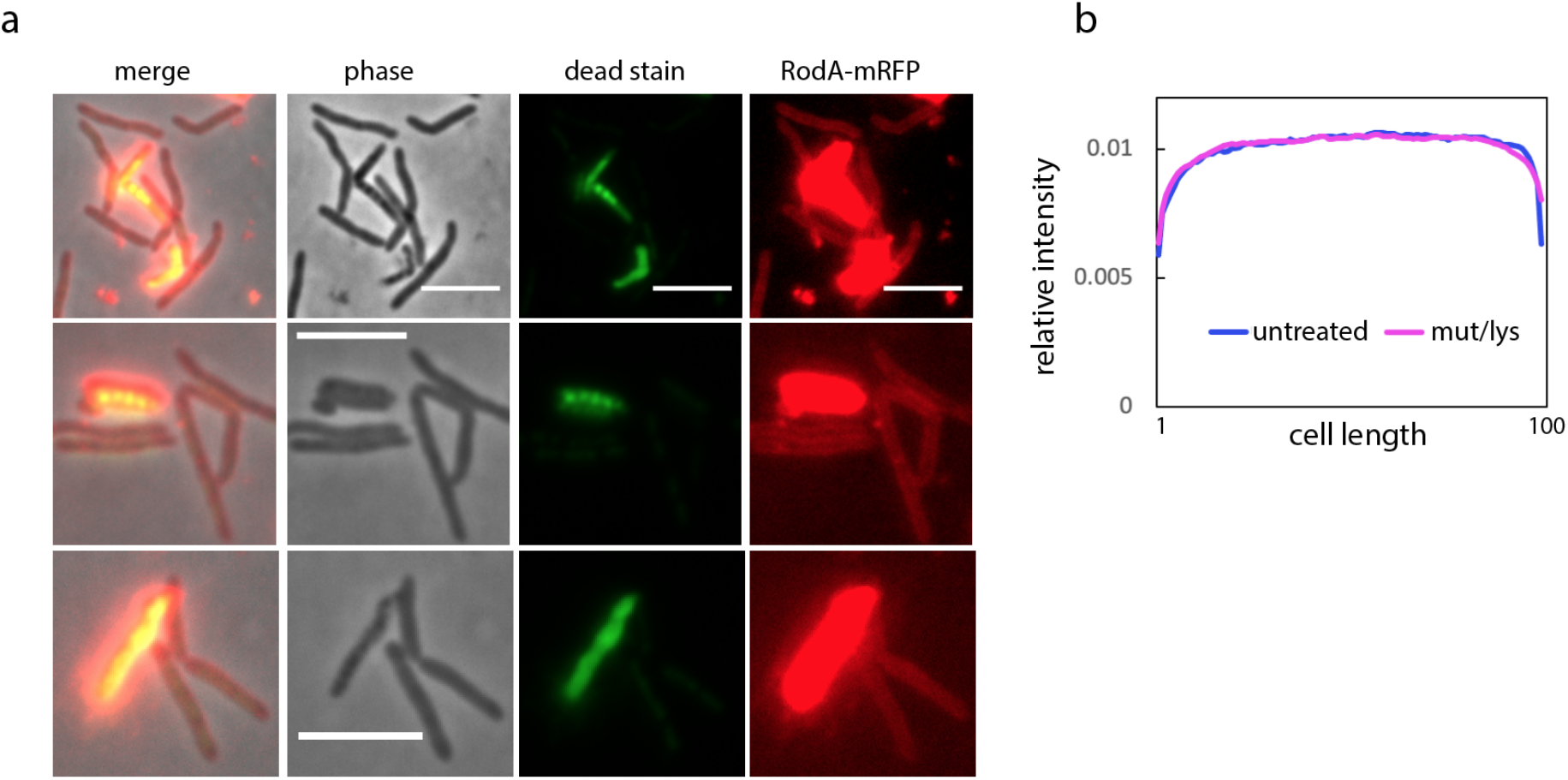
RodA relocalization is not observed in live cells. **(a)** Staining with dead stain SYTOX green reveals that there are no viable cells that display RodA-mRFP polar relocalization phenotype associated with mutanolysin/lysozyme treatment. Scale bars = 5 μm. (b) Cells expressing RodA-mRFP were treated +/− mutanolysin/lysozyme then stained with SYTOX green. Only cells that did not stain green, *i.e*., deemed viable, were included in Oufti followed by MATLAB analysis.

**Figure S7:**
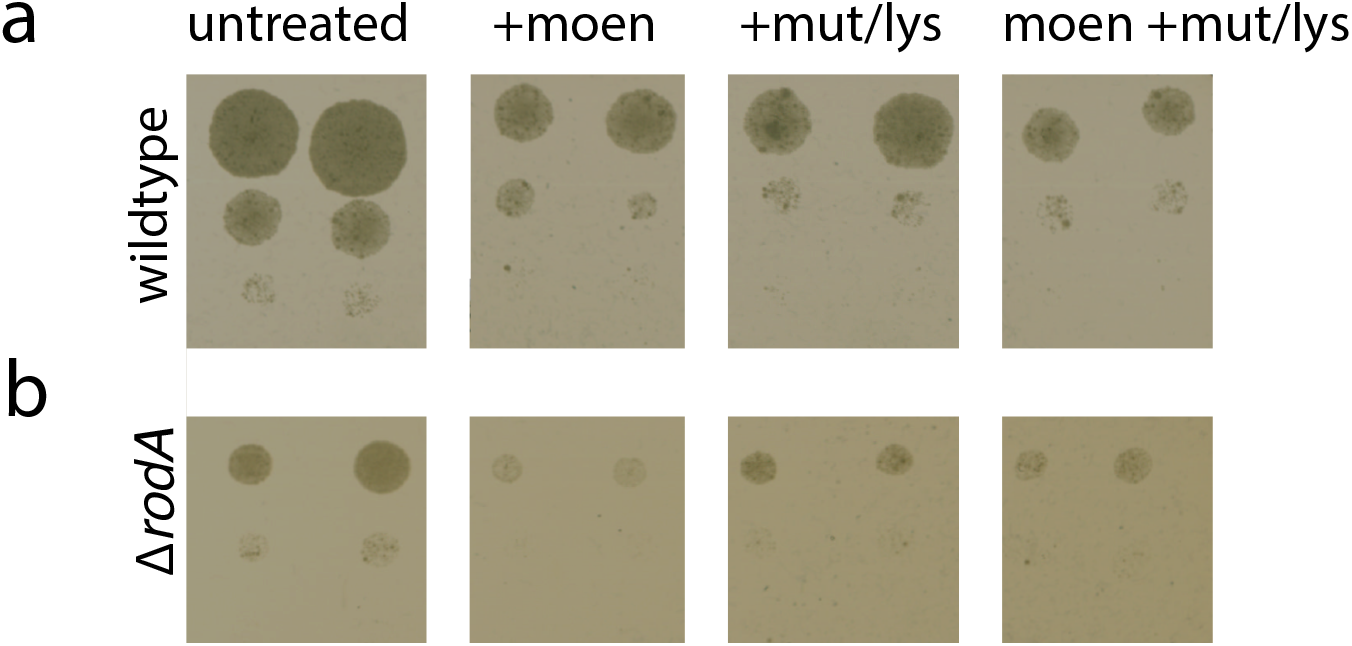
Effect of moenomycin and cell wall digesting enzymes on growth of wildtype and *ΔrodA*. (a) Wildtype and (b) Δ*rodA M. smegmatis* were treated +/− moenomycin (moen) for 30 minutes; mutanolysin/lysozyme (mut/lys) for 60 minutes; or moenomycin followed by mutanolysin/lysozyme. Tenfold serial dilutions were plated following indicated treatment. Moenomycin inhibits growth of wildtype and of Δ*rodA M. smegmatis;* Mutanolysin/lysozyme treatment also inhibits growth in both strains; Combination of moenomycin and mutanolysin/lysozyme treatments does not have an obvious additive effect in either strain. Experiment was performed in triplicate and representative data are shown. a and b not comparable.

**Figure S8:**
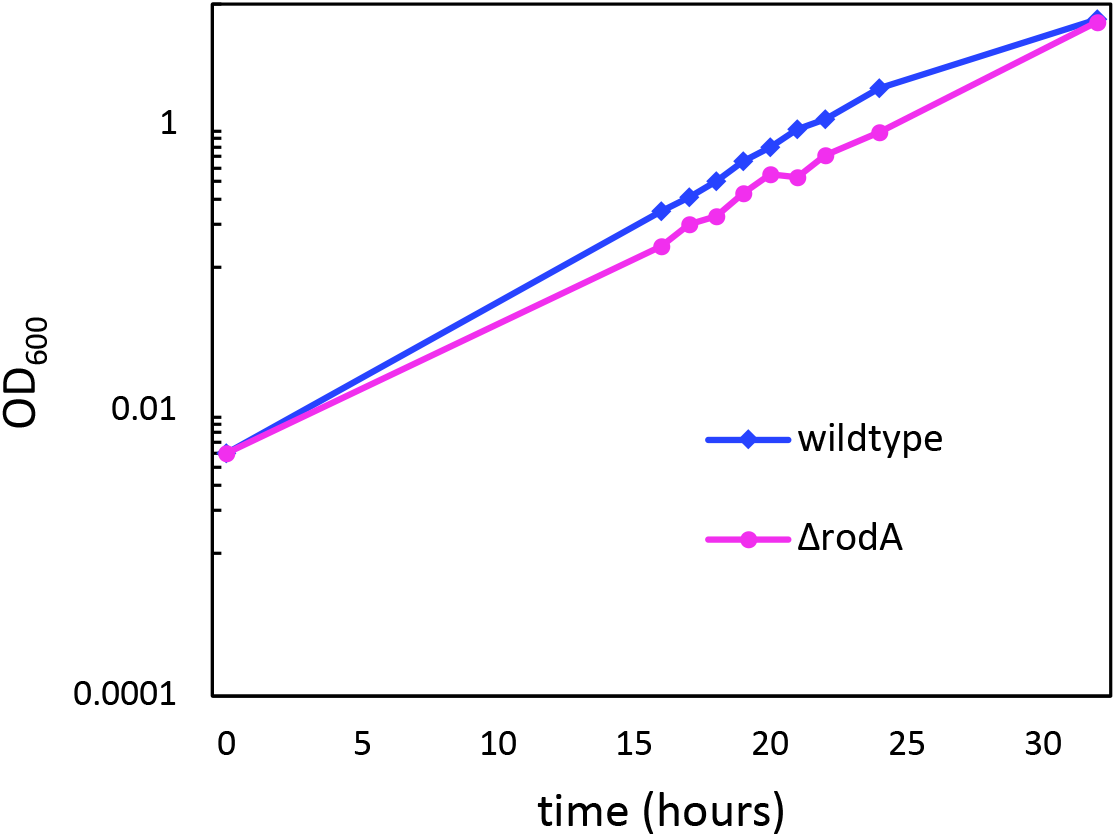
RodA is not required for growth in *M. smegmatis*.

**Table S1:**
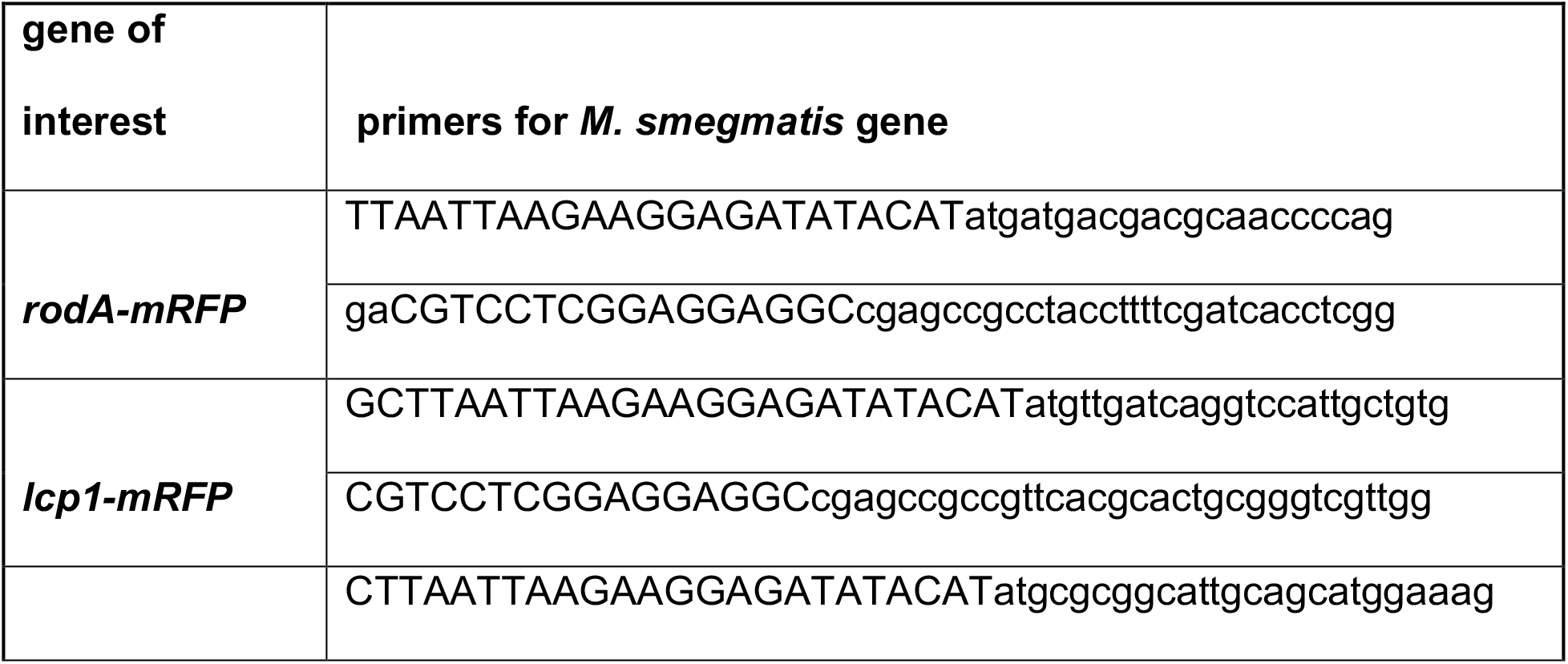
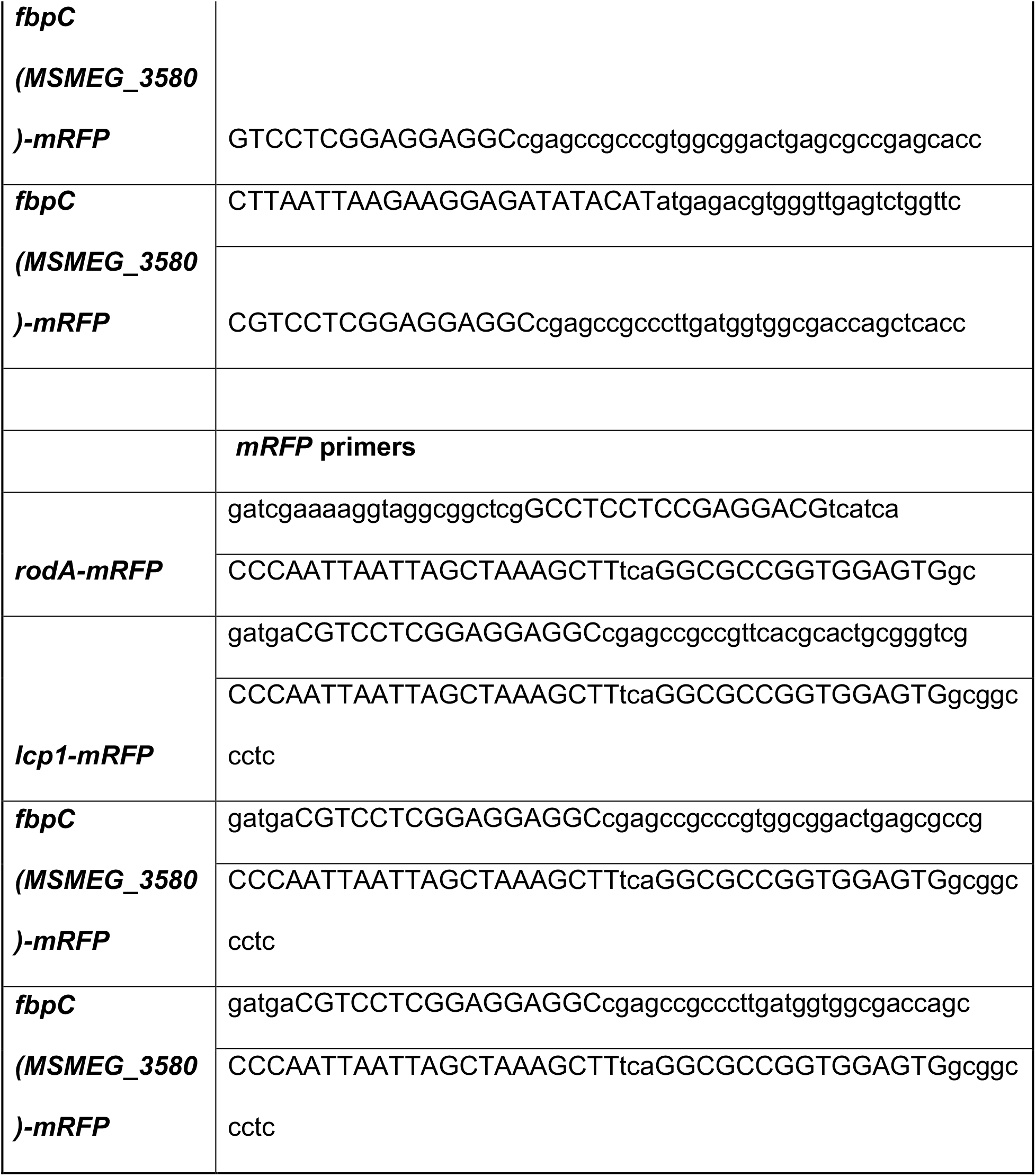
Primers used for fluorescent fusion constructs.

## Notes

Research was supported by funds from the National Institutes of Health (NIH) under awards funding R21 AI144748, R01 AI148255, and DP2 AI138238. ESM was supported by NIH T32 GM008515 administered to the Chemistry Biology Interface Program at the University of Massachusetts Amherst. We are grateful to Ms. Emily Bechtold for technical assistance.

### Competing Interest Statement

The authors have declared no competing interest.

